# Rejuveinix Mitigates Sepsis-Associated Oxidative Stress in the Brain of Mice: Clinical Impact Potential in COVID-19 and Nervous System Disorders

**DOI:** 10.1101/2021.01.03.424883

**Authors:** Fatih M. Uckun, Mehmet Tuzcu, Marcus Gitterle, Michael Volk, Kazim Sahin

## Abstract

Here, we demonstrate that our anti-sepsis and COVID-19 drug candidate Rejuveinix (RJX) substantially improves the survival outcome in the LPS-GalN animal model of sepsis and multi-organ failure. One hundred (100) percent (%) of untreated control mice remained alive throughout the experiment. By comparison, 100% of LPS-GalN injected mice died at a median of 4.6 hours. In contrast to the invariably fatal treatment outcome of vehicle-treated control mice, 40% of mice treated with RJX (n=25) remained alive with a 2.4-fold longer median time survival time of 10.9 hours (Log-rank X^2^=20.60, P<0.0001). Notably, RJX increased the tissue levels of antioxidant enzymes SOD, CAT, and GSH-Px, and reduced oxidative stress in the brain. These findings demonstrate the clinical impact potential of RJX as a neuroprotective COVID-19 and sepsis drug candidate.

## Introduction

Severe viral sepsis caused by SARS-CoV-2, the causative agent of coronavirus disease 2019 (COVID-19), shows a rapid progression associated with a cytokine release syndrome (CRS) and a high case fatality rate due to the development of ARDS and multi-organ failure in high-risk COVID-19 patients (1,2). Our anti-sepsis drug candidate, Rejuveinix (RJX), is a rationally-designed formulation of naturally occurring anti-oxidants and anti-inflammatory compounds with a significant clinical impact potential for COVID-19 associated viral sepsis (3,4). RJX showed a very favorable clinical safety profile in a recently completed double-blind, placebo-controlled, randomized, two-part, ascending dose-escalation Phase 1 study in healthy volunteers (Protocol No. RPI003; ClinicalTrials.gov Identifier: NCT03680105) [18]. It has now entered a randomized double-blind, placebo-controlled Phase 2 study in hospitalized patients with critical COVID-19.

More than a third of patients with COVID-19, especially those with severe to critical COVID-19 who managed on an intensive care unit (ICU) develop central nervous system (CNS) symptoms and signs (e.g., headache, dizziness, ataxia, seizure, delirium, confusion, impaired consciousness) consistent with CNS involvement and /or neurological complications (6–13). Cerebrovascular complications of viral sepsis, including ischemic or hemorrhagic stroke, CNS involvement in CRS, hypoxia as well as the interplay of comorbidities have been implicated as contributing factors (6–13). Due to the neuro-invasive capability of SARS-CoV-2, acute disseminated encephalomyelitis (ADEM) and viral encephalitis have also been suspected in some patients (13).

Oxidative stress caused by the massive production of reactive oxygen species (ROS) is thought to play a major role in the pathogenesis of severe viral sepsis in COVID-19 (14). Chaudry et al. recently proposed that reactive oxygen intermediates and oxidative stress may play an important role in the pathophysiology of COVID-19 associated CNS disease, reminiscent of their role in Parkinson’s disease (15). Notably, RJX exhibited potent anti-oxidant activity and mitigated lipid peroxidation in both prophylactic and therapeutic settings, as reflected by significantly decreased tissue MDA levels and normalization of the tissue levels of the antioxidant enzymes SOD, CAT, and GSH-Px as well as ascorbic acid (5). The primary objective of the present study was to obtain experimental proof of concept that RJX can mitigate sepsis-associated oxidative stress in the brain.

## Materials and Methods

### Rejuveinix (RJX)

RJX is a proprietary composition of naturally occurring anti-oxidants and antiinflammatory agents which, in combination, provide potent and immediate tissue protection. Its ingredients include ascorbic acid, magnesium sulfate heptahydrate, cyanocobalamin, thiamine hydrochloride, riboflavin 5’ phosphate, niacinamide, pyridoxine hydrochloride, and calcium D-pantothenate. RJX is a two-vial system and A and B are each of the two vials. Vial A contains the active ingredients and minerals whereas Vial B contains the buffer, sodium bicarbonate as the Vial A content is acidic (5).

### LPS-GalN Model of Fatal Cytokine Storm, Sepsis, and Multi-organ Failure

The ability of RJX, DEX and RJX plus DEX to prevent fatal shock, ARDS, and multiorgan failure was examined in the well-established LPS-GalN model (5). In this model, LPS is combined with GalN, which further sensitizes mice to LPS-induced systemic inflammatory syndrome and multi-organ failure. The research protocol was approved by the Animal Care and Use Committee of Firat University (Project No. 04052020-391-046). Each mouse received a 500 μL i.p. injection of LPS-GalN (consisting of 100 ng of LPS plus 8 mg of D-galactosamine). Vehicle control mice were treated with 0.5 mL normal saline (NS) intraperitoneally (*i.p*) 2 hours before and 2 hours after the *i.p* injection of LPS-GalN. Test mice received 0.7 mL/kg RJX dose (2 hours before and 2 hours after LPS-GalN. Mice were monitored for mortality for 24 h. The Kaplan–Meier method, log-rank chi-square test, was used to analyze the 24 h survival outcomes of mice in the different treatment groups. At the time of death, brains were harvested for biochemical studies.

Lipid peroxidation as a biomarker of oxidative stress was determined and expressed as the amount of malondialdehyde (MDA, nmol/g tissue) in the brain, as previously described (5). The enzymatic activities of SOD, CAT and GSH-Px in the brain specimens were determined using the commercially available kits (Cayman Chemical, Ann Arbor, MI, USA) according to the manufacturer’s procedures (5).

### Statistical Analyses

Statistical analyses employed standard methods, including analysis of variance (ANOVA) and/or, nonparametric analysis of variance (Kruskall-Wallis) using the SPSS statistical program (IBM, SPPS Version 21), as reported (5). Furthermore, the Kaplan-Meier method, log-rank chi-square test, was used to investigate survival and fatality in each group. P-values <0.05 were considered significant.

## Results

### RJX improves survival outcome after LPS-GalN induced sepsis

One hundred (100) percent (%) of untreated control mice remained alive throughout the experiment. By comparison, 100% of LPS-GalN injected mice died at a median of 4.6 hours (**Figure 1**). RJX was examined for its protective activity at a dose level, which is >10-fold lower than its maximum tolerated dose (MTD) of 0.759 mL/kg for human subjects (viz.; 4.2 mL/kg of a 6-fold diluted solution). RJX-treated mice had an improved survival outcome after being injected with LPS-GalN. In contrast to the invariably fatal treatment outcome of vehicle-treated control mice, 40% of mice treated with RJX (n=25) remained alive with a 2.4-fold longer median time survival time of 10.9 hours (Log-rank X^2^=20.60, P<0.0001) (**Figure 1**).

**Figure 1.**
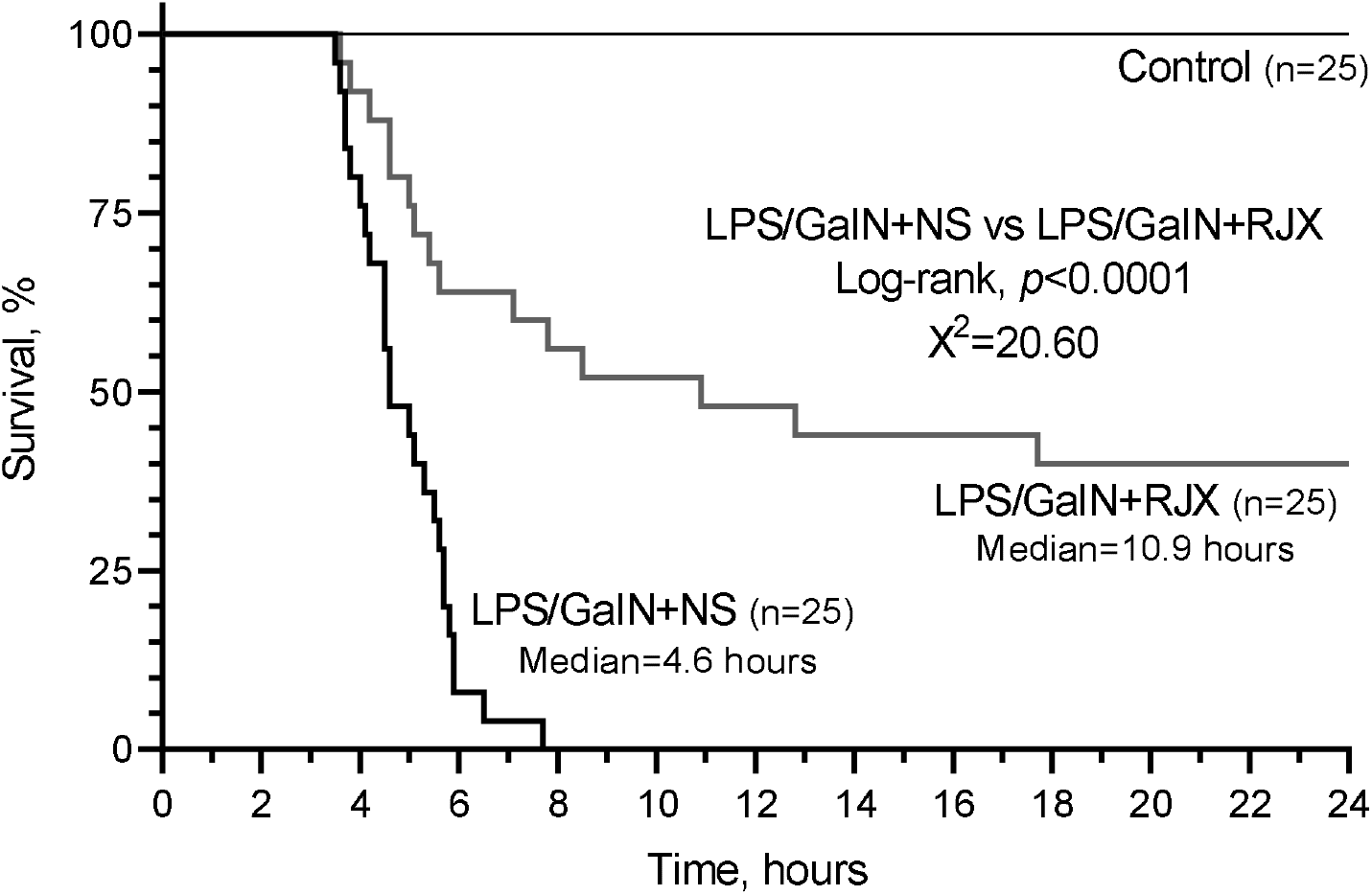
In Vivo Protective Activity of RJX in the LPS-GALN challenged mice. Groups of 25 BALB/C mice were treated with i.p injections of 6-fold diluted RJX (4.2 mL/kg, 0.5 ml/mouse) or vehicle (NS) 2 hours before or post-injection of LPS-GalN. Except for untreated mice (Control), each mouse received 0.5 ml of LPS-GalN (consisting of 100 ng of LPS plus 8 mg of D-galactosamine) i.p. Survival is shown as a function of time after the LPS-GalN challenge. Depicted are the survival curves for each group along with the median survival times and the log-rank P-value for the comparison of the LPS-GalN+RJX group versus the LPS/GaIN+NS group.

### RJX reduces the oxidative stress in the brain after LPS-GalN induced sepsis

No histopathologic brain lesions were observed in any of the mice challenged with LPS-GalN. However, the brain MDA levels measuring lipid peroxidation were markedly elevated, and the levels of the antioxidant enzymes SOD, CAT, and GSH-Px in the brain were markedly reduced in LPS-GalN treated mice consistent with severe oxidative stress (**Figure 2**). RJX decreased the brain MDA levels and normalized in a dose-dependent manner the reduced levels of the antioxidant enzymes SOD, CAT, and GSH-Px.

**Figure 2.**
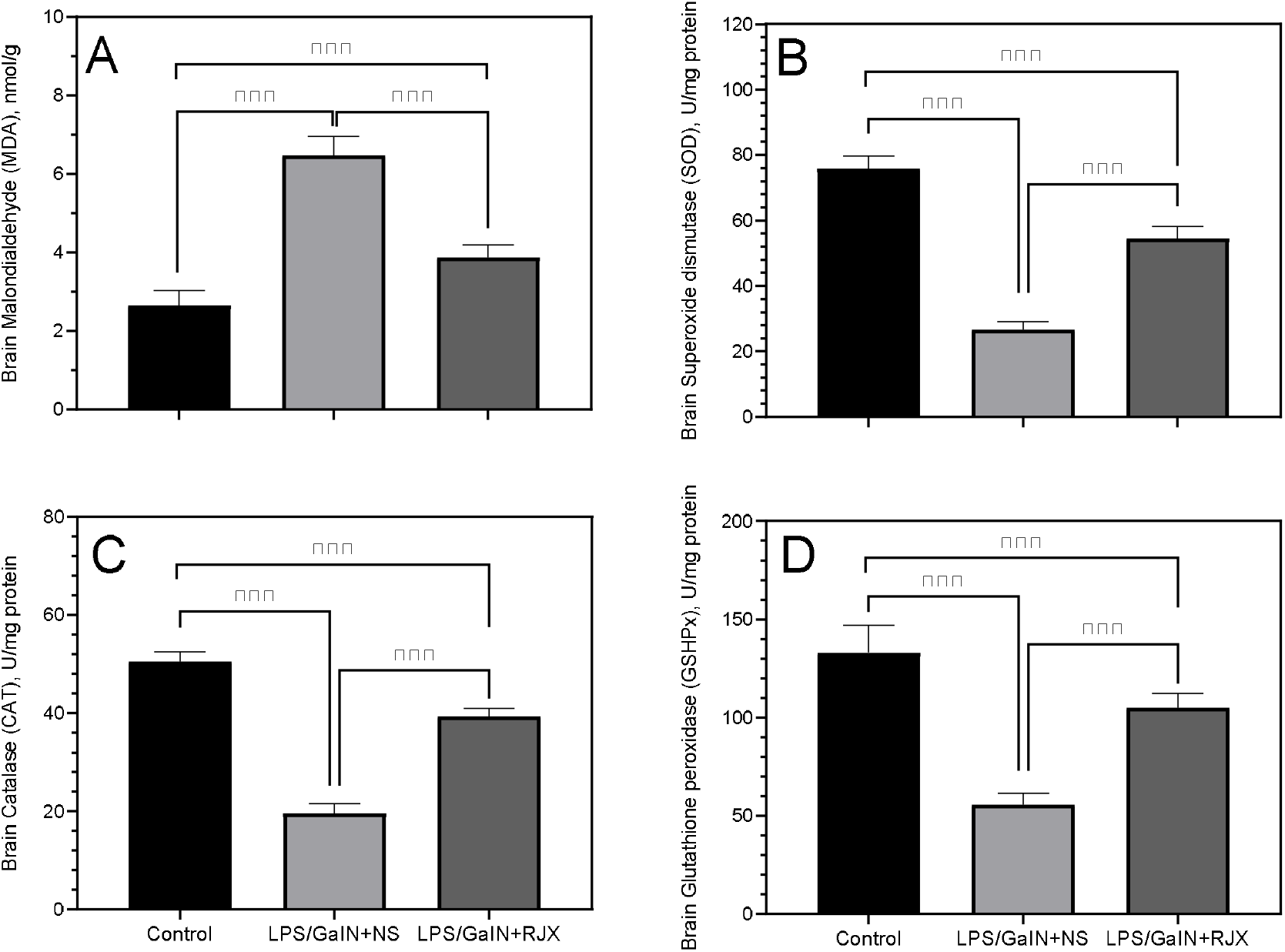
Effect of Rejuveinix (RJX) on brain malondialdehyde (MDA; Panel A), superoxide dismutase (SOD; Panel B), catalase (CAT; Panel C) and Glutathione peroxidase (GSHPx; Panel D) in the lipopolysaccharide-galactosamine (LPS-GalN) challenged mice. Each bar represents the mean and standard deviation for the measured parameter of mice from each specific treatment group (6 mice/group). BALB/C mice were treated with i.p injections of 6-fold diluted RJX (4.2 mL/kg, 0.5 ml/mouse) or vehicle (NS) 2 hours before or post-injection of LPS-GalN. Except for untreated mice (Control), each mouse received 0.5 ml of LPS-GalN (consisting of 100 ng of LPS plus 8 mg of D-galactosamine) i.p. (ANOVA and Tukey’s post-hoc test. Statistical significance between groups is shown by: *** P < 0.001).

## Discussion

Patients with high-risk COVID-19 are in urgent need of treatment platforms capable of preventing the disease progression and/or reducing the case mortality rate by stopping or reversing the pulmonary as well as systemic inflammatory process that cause the ARDS and culminate in multi-organ failure (1,2,16). Here we extend our previous study and provide experimental evidence that RJX is capable of improving the survival outcome in the LPS-GalN mouse model of fatal sepsis.

SOD, CAT, and GSH-Px are three pivotal antioxidant defense enzymes, and their levels are altered by the level of oxidative stress that is a hallmark of severe inflammation of sepsis (14). Oxidative stress caused by massive production of reactive oxygen species (ROS) is thought to play a major role in the pathogenesis of severe viral sepsis in COVID-19 as well (14). The MDA levels were markedly elevated in the brain specimens of the LPS-GalN challenged mice which is consistent with increased lipid peroxidation. In parallel, the levels of the antioxidant enzymes SOD, CAT, and GSH-Px were profoundly suppressed due to severe oxidative stress. RJX exhibited potent anti-oxidant activity and mitigated lipid peroxidation, as reflected by significantly decreased tissue MDA levels and normalization of the tissue levels of the antioxidant enzymes SOD, CAT, and GSH-Px as well as ascorbic acid. We hypothesize that RJX will shorten the time to resolution of ARDS and viral sepsis in COVID-19 patients by preventing the development of a fulminant cytokine storm as well as reversing the cytokine-mediated multi-system inflammatory process and oxidative stress, thereby mitigating the inflammatory organ injury.

Oxidative stress owing to mitochondrial dysfunction has been implicated in the pathophysiology of Alzheimer’s disease (AD), Parkinson’s disease (PD), and Huntington’s disease (HD). Notably, in a mouse CNS model of severe oxidative stress, RJX rapidly and substantially increased the levels of the anti-oxidant enzymes SOD, CAT, GSH-Px that were reduced in the brains of LPS-GalN treated mice consistent with severe oxidative stress. Our results indicate that RJX could have therapeutic utility in the treatment of AD, PD, HD, and amyotrophic lateral sclerosis (ALS). Furthermore, because of the well-established role of oxidative stress in the development and progression of ischemic stroke, RJX could significantly improve the standard of care for stroke as well.

## Author contributions

Each author has made significant and substantive contributions to the study, reviewed and revised the manuscript, provided final approval for submission of the final version. No medical writer was involved. F.M.U conceived the study, designed the evaluations reported in this paper, directed the data compilation and analysis, analyzed the data, and prepared the initial draft of the manuscript. Each author had access to the source data used in the analyses.

